# Magnesium suppresses defects in the formation of 70S ribosomes as well as in sporulation caused by lack of several individual ribosomal proteins

**DOI:** 10.1101/300103

**Authors:** Genki Akanuma, Kotaro Yamazaki, Yuma Yagishi, Yuka Iizuka, Morio Ishizuka, Fujio Kawamura, Yasuyuki Kato-Yamada

## Abstract

Individually, the ribosomal proteins L1, L23, L36 and S6 are not essential for cell proliferation of *B. subtilis*, but the absence of any one of these ribosomal proteins causes a defect in the formation of the 70S ribosomes and a reduced growth rate. In mutant strains individually lacking these ribosomal proteins, the cellular Mg^2+^ content was significantly reduced. The deletion of YhdP, an exporter of Mg^2+^, and overexpression of MgtE, the main importer of Mg^2+^, increased the cellular Mg^2+^ content and restored the formation of 70S ribosomes in these mutants. The increase in the cellular Mg^2+^ content improved the growth rate of the Δ*rplA* (L1) and the Δ*rplW* (L23) mutant but did not restore those of the Δ*rpmJ* (L36) and the Δ*rpsF* (S6) mutants. The lack of L1 caused a decrease in the production of Spo0A, the master regulator of sporulation, resulting in a decreased sporulation frequency. However, deletion of *yhdP* and overexpression of *mgtE* increased the production of Spo0A and partially restored the sporulation frequency in the Δ*rplA* (L1) mutant. These results indicate that Mg^2+^ can partly complement the function of several ribosomal proteins, probably by stabilizing the conformation of the ribosome.

**IMPORTANCE:** We previously reported that an increase in the cellular Mg^2+^ content can suppress defects in 70S ribosome formation and growth rate caused by the absence of ribosomal protein L34. In the present study, we demonstrated that even in mutants lacking individual ribosomal proteins other than L34 (L1, L23, L36 and S6), an increase in the cellular Mg^2+^ content could restore the 70S ribosome formation. Moreover, the defect in sporulation caused by the absence of L1 was also suppressed by an increase in the cellular Mg^2+^ content. These findings indicate that at least part of the function of these ribosomal proteins can be complemented by Mg^2+^, which is essential for all living cells.

## INTRODUCTION

The bacterial ribosome (70S), which plays a central role in protein synthesis, is a complex macromolecule that is composed of small (30S) subunit and large (50S) subunits. The small subunit is comprised of the 16S rRNA and more than 20 proteins, whereas the large subunit is comprised of the 23S and 5S rRNAs and more than 30 proteins (1, 2). Protein synthesis by the ribosome, called translation, requires the coordinated action of these subunits. The small subunit associates with the mRNA and the anticodon stem-loop of the bound tRNA, and engages in ensuring the fidelity of translation by checking for correct pairing between the codon and anticodon (3–7). The large subunit associates with the acceptor arms of the tRNA and catalyzes the formation of a peptide bond between the amino acid attached to the tRNA in the A-site and the nascent peptide chain bound to the tRNA in the P-site (8, 9). The ribosomal proteins that constitute these subunits play important role(s) in translation. For instance, ribosomal protein L1, which is localized to the stalk region near the E-site (10, 11), plays a critical role in the translocation of the newly deacylated tRNA from the P to the E site (12). Ribosomal protein L2 plays important roles in binding of the tRNA to the A and P sites, peptidyltransferase activity, and formation of the peptide bond (13–17). Therefore, the mature conformation of the 70S ribosomes is required for efficient translation activity. Although the ribosomal proteins are important in the translation processes as well as in the association of the ribosomal subunits (13, 18, 19), several genes encoding ribosomal protein can be deleted. In *Escherichia coli*, 22 of the 54 genes for ribosomal proteins are not individually essential for cell proliferation (20, 21). Similarly, in *Bacillus subtilis*, 22 of the 57 genes for ribosomal proteins can be individually deleted (22). The *rpmH* gene, encoding ribosomal protein L34, which is a component of the large subunit, is one of the nonessential genes. Mutants lacking L34 have a severe defect in the formation of the 70S ribosome and a reduced growth rate (22). However, we found that the defect in the formation of 70S ribosomes and the reduction in the growth late could be suppressed by an increase in the Mg^2+^ content in the cell (23). Magnesium ions are the most abundant divalent cations in living cells (24, 25), and are important for the maintenance of ribosome structure. Mg^2+^ is required for both stabilization of the secondary structure of rRNA and binding of the ribosomal proteins to the rRNA (26–28). The *in vitro* association of the 30S and 50S ribosomal subunits to form intact 70S ribosomes depends strongly on the concentration of Mg^2+^ (29–31). Therefore, we believe that Mg^2+^ can partly complement the L34 function by stabilizing both the conformation of the 50S subunit and the intersubunit bridges.

In the present study, to elucidate whether Mg^2+^ can also complement mutant strains lacking ribosomal proteins other than L34, we examined the effect of increasing the Mg^2+^ content in mutant strains individually lacking ribosomal proteins L1, L23, L36 and S6 on the formation of 70S ribosomes, the growth rate, and on sporulation.

## RESULTS

### Reduction in the cellular Mg^2+^ content caused by lack of ribosomal proteins was restored by disruption of *yhdP* and overexpression of *mgtE*

The defect in the formation of 70S ribosomes caused by the absence of L34 could be suppressed by increasing the cellular Mg^2+^ content (23). To investigate the generality of the partial complementation of the ribosomal-protein function by Mg^2+^, a disruption of *yhdP* and the multicopy plasmid pDGmgtE, which can induce the overexpression of *mgtE*, were introduced into mutants lacking individual ribosomal proteins L1, L23, L36 and S6. MgtE is the main importer of Mg^2+^ (32), whereas YhdP is probably an exporter of Mg^2+^ in *B. subtilis* (23). We previously reported that the absence of L34 (RpmH) caused a decrease in the Mg^2+^ content in the cell, probably due to a reduced number of 70S ribosomes, and that the Mg^2+^content in the Δ*rpmH* mutant was restored by disruption of *yhdP* and overexpression of *mgtE* (23). Similarly, the Mg^2+^ contents in the Δ*rplA* (L1), Δ*rplW* (L23) and Δ*rpmJ* (L36) mutants were also significantly reduced (Fig. 1). However, the Mg^2+^ content in these three mutants was restored, albeit incompletely, by disruption of *yhdP* and overexpression of *mgtE* (Fig. 1). In this experiment, the cellular Mg^2+^ concentration was calculated by dividing the amount of Mg^2+^ per cell by the cell volume. The cell volume of each mutant was estimated from the cell size, which was measured by microscopic analysis, as described in Materials and Methods. However, the cell size of the Δ*rpsF* (S6) mutant could not be defined, because the cellular morphology of the Δ*rpsF* mutant was aberrantly filamentous (Fig. S1). Thus, in the Δ*rpsF* mutant, the relative Mg^2+^amount per cell, when the Mg^2+^amount of a cell in the parental strain was defined as 1, is shown in Fig. 1. Although a comparison of the Mg^2+^ content in the Δ*rpsF* mutants with that in the wild type was difficult, the Mg^2+^content in the Δ*rpsF* mutants was certainly increased by disruption of *yhdP* and overexpression of *mgtE.* It should be noted that the Mg^2+^ ions that were chelated in ribosomes and other enzymes were also detected by this method, because the cells were completely disrupted by sonication and proteins were denatured by acid treatment.

**Fig. 1.**
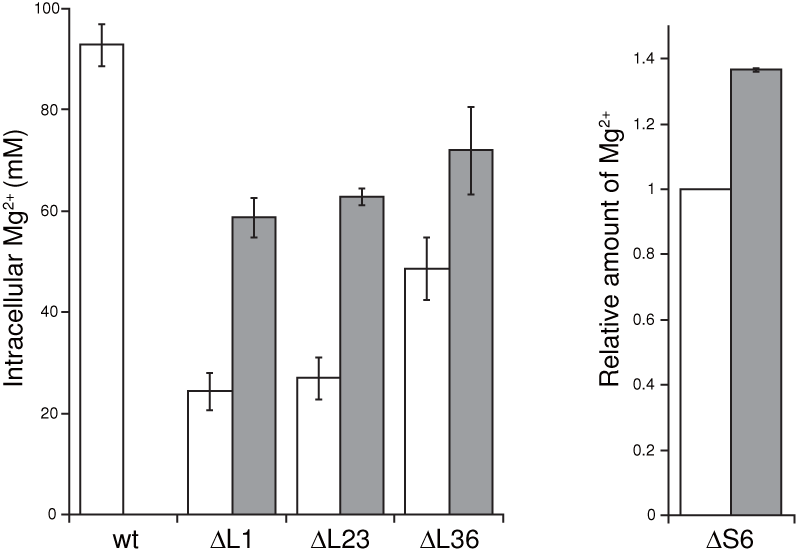
Reduction in the Mg^2+^ content in mutant strains lacking individual ribosomal proteins and partial restoration by disruption of*yhdP* and overexpression of *mgtE.* The Mg^2+^ content per cell in exponential phase, which was measured as described in the Materials and Methods, is shown. In the case of the Δ*rpsF* (S6) mutant, the relative amount of Mg^2+^ per cell is shown (see text for details). White bars indicate the wild type and each mutant lacking individual ribosomal proteins. Gray bars indicate the results when *yhdP* was disrupted and *mgtE* was overexpressed. The means of three independent experiments are shown. Error bars indicate standard deviations.

### The effect of increasing the cellular Mg^2+^ content of mutants lacking individual ribosomal proteins on the formation of 70S ribosomes and the growth rate

As shown in Fig. 2, the lack of individual ribosomal proteins (L1, L23, L36, S6) caused defects in the formation of 70S ribosomes that are consistent with our previous data (22). The defect in 70S-ribosome formation observed in these mutants was suppressed by disruption of *yhdP* and overexpression of *mgtE*, to varying degrees (Fig. 2). In all of the mutants investigated here, the amount of 70S ribosomes relative to the amount of dissociated subunits was restored by increasing the Mg^2+^ content in the cell. These results indicate that Mg^2+^ can suppress the defect in the formation of 70S ribosomes caused by the absence of several individual ribosomal proteins.

**Fig. 2.**
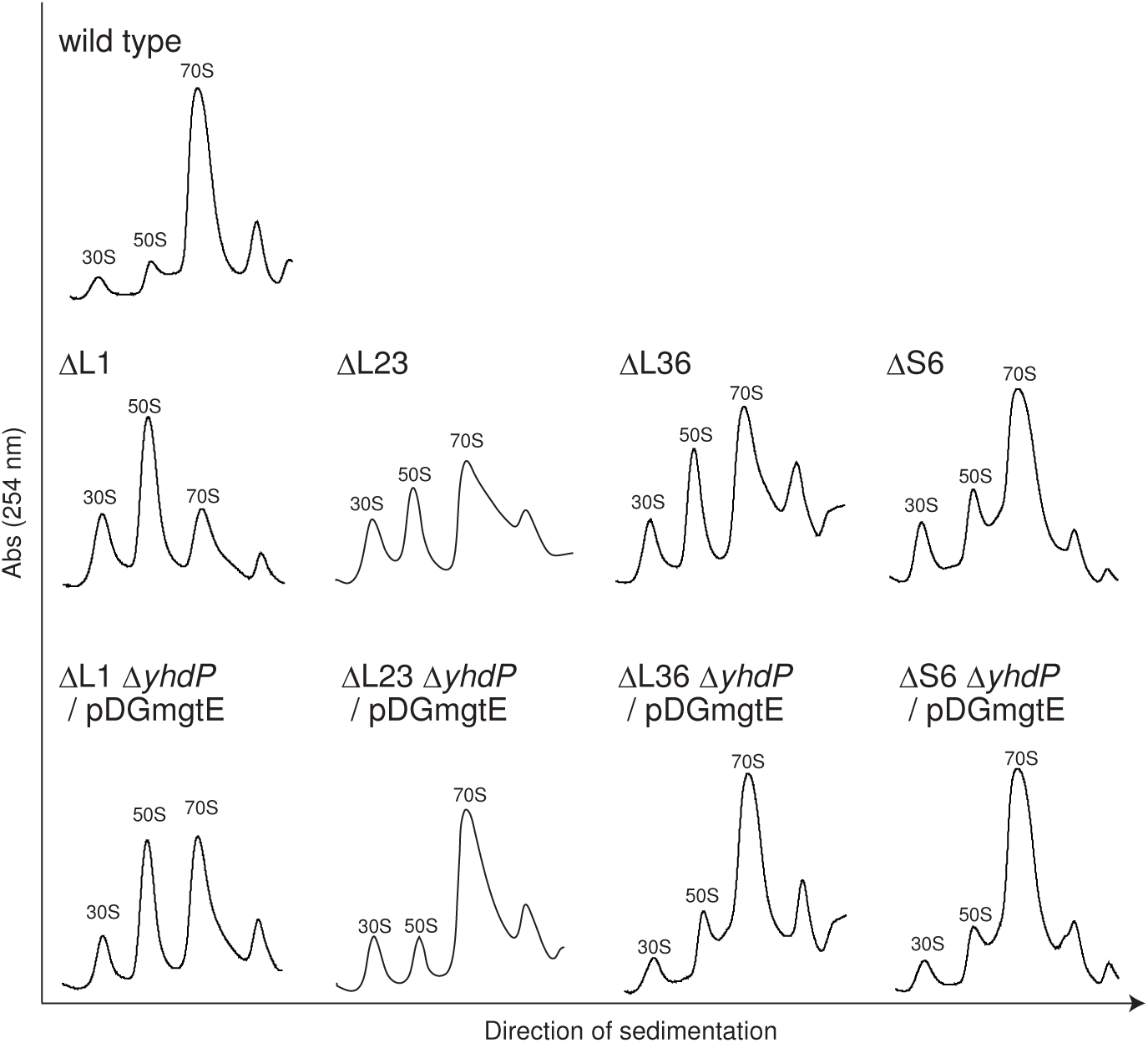
Defect in 70S ribosome formation in the absence of each ribosomal protein and its suppression by the disruption of *yhdP* and overexpression of *mgtE.* Crude cell extracts were sedimented through a 10–40% sucrose gradient as described in the Materials and Methods. The 30S, 50S, and 70S peaks are indicated in each individual profile. The term ‘/ pDGmgtE’ indicates the overexpression of *mgtE* in the mutant cells.

We next investigated the effect of the cellular Mg^2+^ content on the growth rate of the mutants. We have reported that the slow growth observed in the Δ*rpmH* (L34) mutant was suppressed by an increase in the Mg^2+^ content, probably due to the restoration of the amount of 70S ribosomes (23). A reduction in the growth rate was observed in the mutants lacking individual ribosomal proteins, which agrees with our previous results (22). As expected, in the Δ*rplA* (L1) and Δ*rplW* (L23) mutants, the growth rate was partially restored by disruption of *yhdP* and overexpression of *mgtE* (Fig. 3A and B, Table 1). When only *mgtE* was overexpressed in the Δ*rplA* (L1) mutant, its effect on the growth rate was minimal (23). The combination of overexpression of *mgtE* and disruption of *yhdP*, however, increased the growth rate of the Δ*rplA* (L1) mutant. In contrast, the growth rates of the Δ*rpmJ* (L36) and Δ*rpsF* (S6) mutants were not significantly increased when the cellular Mg^2+^ content was increased (Fig. 3C and D, Table 1). Therefore, the increased formation of 70S ribosomes did not necessarily restore the growth rate of the mutants lacking individual ribosomal proteins.

**Fig. 3.**
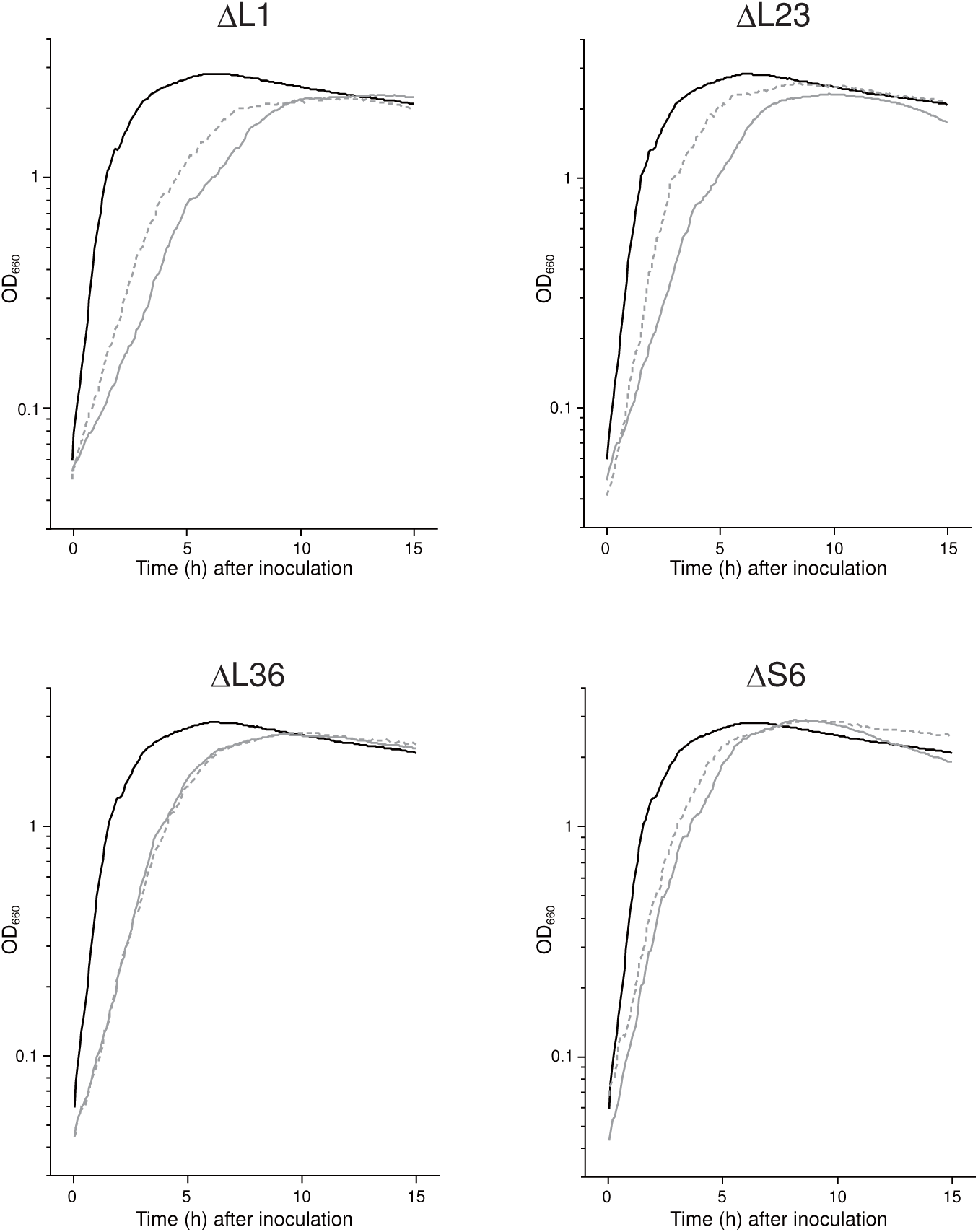
Effects of the increase in cellular Mg^2+^ content on the growth rate of the mutant strains lacking individual ribosomal proteins. Cells were grown in LB at 37°C, and the optical density at 660 nm was measured. Growth curves of wild type and each mutant lacking individual ribosomal proteins are shown using black solid lines and gray solid lines, respectively. Growth curves of each mutant in which *yhdP* was disrupted and *mgtE* was overexpressed are shown by gray dotted lines.

**Table 1.** Doubling times (min) of mutants lacking ribosomal proteins.

| Strain | Parental | $\Delta yhdP/pDGmgtE$ |
| --- | --- | --- |
| wt | $21.4 \pm 1.4$ | |
| $\Delta rplA$ (L1) | $67.7 \pm 1.7$ | $54.6 \pm 0.49$ |
| $\Delta rplW$ (L23) | $56.7 \pm 1.9$ | $37.0 \pm 2.4$ |
| $\Delta rpmJ$ (L36) | $43.3 \pm 1.3$ | $41.2 \pm 0.54$ |
| $\Delta rpsF$ (S6) | $38.2 \pm 1.1$ | $36.4 \pm 0.49$ |

Means of three independent experiments with ± SD are shown.

### The increase in the cellular Mg^2+^content suppresses the defect in sporulation caused by the absence of ribosomal protein L1

We previously found that the absence of ribosomal protein L1 causes a defect in sporulation (22). It should be note that this phenotype was not caused solely by the decreased growth rate, because the sporulation frequency of the Δ*rpmH* mutant, which also showed a severe growth defect similar to that of the Δ*rplA* (L1) mutant, was almost the same as that of the wild type (22). We therefore investigated whether the sporulation defect of the Δ*rplA* (L1) mutant could be suppressed by Mg^2+^. Consistent with our previous data, the Δ*rplA* (L1) mutant was severely defective in forming heat-resistant spores (the sporulation frequency was less than 0.01%) (Table 2). However, the sporulation frequency of the Δ*rplA* (L1) mutant was significantly restored by disruption of *yhdP* and overexpression of *mgtE* (Table 2). In addition, the growth rate of the Δ*rplA* (L1) mutant in sporulation medium was also restored by disruption of *yhdP* and overexpression of *mgtE* (Fig. 4A). These results indicate that increasing the cellular Mg^2+^ content can suppress not only the growth defect, but also the sporulation defect in the Δ*rplA* (L1) mutant.

**Table 2.** Restoration of sporulation frequency of the ΔL1 mutant by Mg^2+^

| Strain | C.f.u. ml<sup>-1</sup>* |  | Frequency (%)<sup>*</sup> |
| --- | --- | --- | --- |
|  | Total | Spores |  |
| wt | $6.4 \times 10^8$ | $5.8 \times 10^8$ | $90 \pm 7.9$ |
| $\Delta L1$ | $1.6 \times 10^8$ | $2.3 \times 10^2$ | $1.7 (\pm 1.2) \times 10^{-4}$ |
| $\Delta L1 \Delta yhdP$ / pDGmgtE | $3.1 \times 10^8$ | $7.5 \times 10^7$ | $25 \pm 4.7$ |

\*Means of three independent experiments ( ± SD for sporulation frequency).

**Fig. 4.**
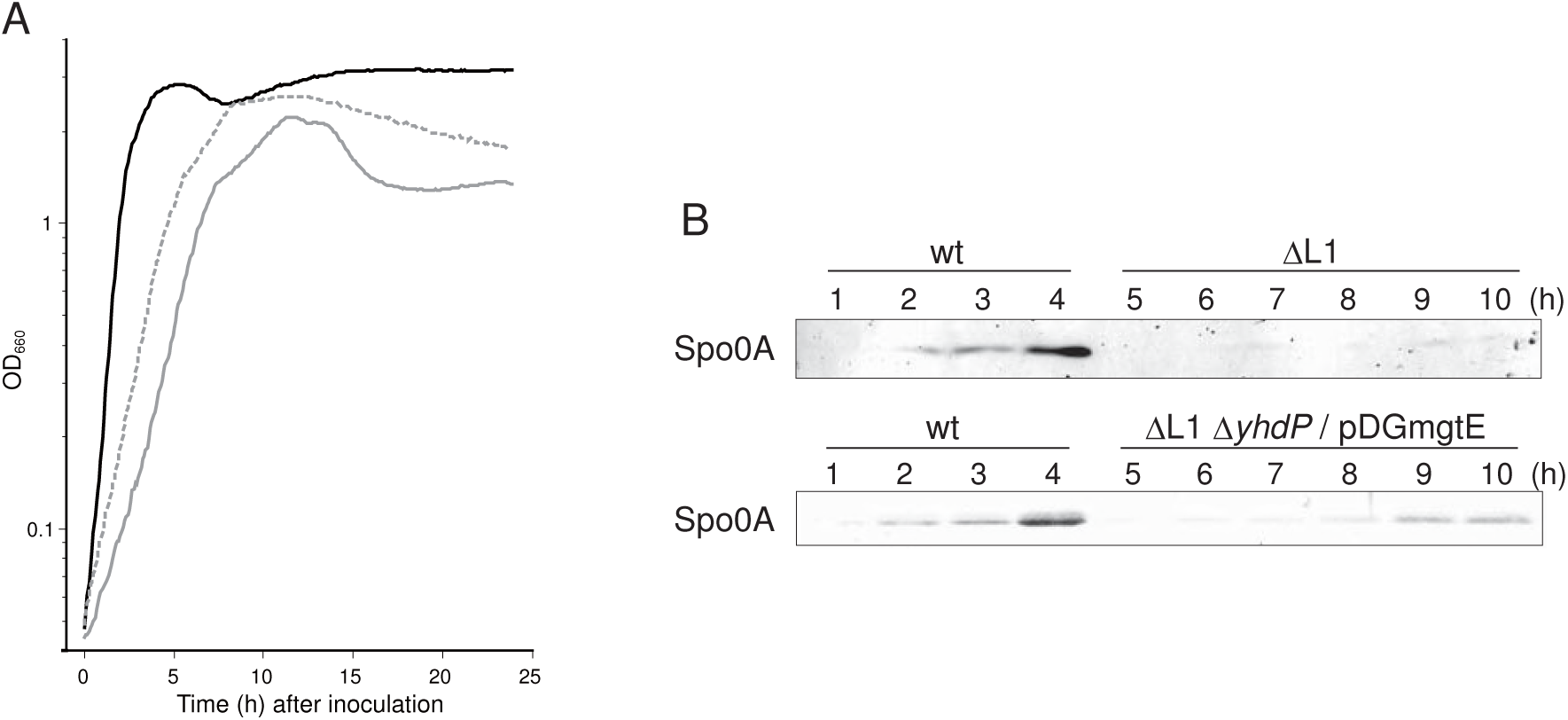
Reduction in the growth rate and in the production of Spo0A caused by lack of L1 and their suppression by the disruption of *yhdP* and overexpression of *mgtE.* (A) Cells were grown in sporulation medium (2×SG) at 37°C, and the optical density at 660 nm was measured. Growth curves of wild type and the Δ*rplA* (L1) mutant are shown using black and gray solid lines, respectively. The growth curve of the L1 mutant in which *yhdP* was disrupted and *mgtE* was overexpressed is shown by a gray dotted line. (B) Cells were grown in sporulation medium at 37°C, and were collected at the indicated times. Crude cell extracts were subjected to western blot analysis using antisera against the Spo0A. The term ‘/ pDGmgtE’ indicates the overexpression of *mgtE* in the mutant cells.

The restoration of spore formation by the Δ*rplA* (L1) mutant prompted us to identify which stage of sporulation was affected by the absence of L1 and restoration by Mg^2+^. At the initiation stage of *B. subtilis* sporulation, cells divide asymmetrically, and chromosomal DNA is concentrated in the forespore (33). In fact, asymmetric septation and concentration of chromosomal DNA were detected in the wild-type cells five h after inoculation in sporulation medium (Fig. S2). In contrast, in the Δ*rplA* (L1) cells, asymmetric septation was not observed even 24 h after inoculation (Fig. S2). However, the disruption of *yhdP* and overexpression of *mgtE* helped formation of the asymmetric septum in the Δ*rplA* (L1) cells (Fig. S2). We next examined the level of Spo0A in the Δ*rplA* (L1) mutant. Phosphorylation of Spo0A, the master transcriptional regulator of sporulation, governs the decision to initiate sporulation (34–36).In wild-type cells, the level of Spo0A increased four hours after inoculation into sporulation medium, whereas Spo0A was barely detectible in Δ*rplA* (L1) cells even 10 h after inoculation (Fig. 4B). The disruption of *yhdP* and overexpression of *mgtE* in the Δ*rplA* (L1) cells increased the amount of Spo0A by 9 h after inoculation, although the level of Spo0A remained lower than that in wild type (Fig. 4B). These results indicate that the defect in the initiation stage of sporulation caused by the absence of L1 can be at least partially suppressed by an increase in the Mg^2+^ content in the cell.

## DISCUSSION

The cellular Mg^2+^ contents of the mutant strains individually lacking L1, L23, or L36 were reduced compared to that of wild type, while that of the Δ*rpsF* (S6) mutant was difficult to calculate because of its filamentous cellular morphology (Fig. 1 and Fig. S1). The reduction in the amount of Mg^2+^ was probably caused by the lower amount of 70S ribosomes, which harbor more than 170 Mg^2+^ ions per complex (37), and by a decrease in the amount of protein and RNA other than ribosomes that can chelate Mg^2+^. In fact, we previously showed that the reduction in the cellular Mg^2+^ content correlated with the decrease in the amount of 70S ribosomes (23). On the other hand, the disruption of *yhdP* and overexpression of *mgtE* increased cellular Mg^2+^ content and restored the formation of 70S ribosomes in the mutants lacking individual ribosomal proteins tested here (Fig. 2). Although the absence of L34 causes ribosomal protein L16 to dissociate from the 50S subunit, the increase in the Mg^2+^ content restores the binding of L16 to the 50S subunit, indicating that Mg^2+^ can stabilize the conformation of 50S subunits lacking L34 (23). Likewise, stabilization of the conformation of each subunit as well as inter bridges between the subunits by Mg^2+^ probably restored 70S formation in the mutants lacking individual ribosomal proteins tested here (L1, L23, L36 and S6).

The increase in the 70S ribosome formation restored cellular translational activity that may have resulted in the suppression of the defect in the growth rate of the Δ*rplA* (L1) and Δ*rplW* (L23) mutants (Fig. 3 and Table 1). However, the restoration of growth rate of these mutants was only partial. Possible reasons for this partial restoration of the growth are (i) an incomplete restoration of the normal amount of 70S ribosomes, and (ii) functions of the ribosomal proteins other than in stabilizing the 70S ribosomes could not be complemented by Mg^2+^. Ribosomal protein L1 plays a critical role in the translocation of the newly deacylated tRNA from the P to the E site (12), while ribosomal protein L23, which is located at the polypeptide exit channel of the large subunit, tethers trigger factor to the ribosome (38). Trigger factor, which is the first molecular chaperone interacting with newly synthesized polypeptides by the ribosome, promotes protein folding (39–41). The functions of these ribosomal proteins are probably essential for efficient growth. In contrast to mutants lacking L1 or L23, the growth rates of the Δ*rpmJ* (L36) and Δ*rpsF* (S6) mutants were not significantly increased when the Mg^2+^ content was increased, although 70S ribosome formation was restored to near wild-type levels (Fig. 2 and Fig. 3). Although the detailed functions of L36 and S6 in protein synthesis are unknown, their role(s) in translation or other function(s) do not appear to be complemented by Mg^2+^. In addition, the filamentous morphology of cells caused by the absence of S6 was not repaired by increasing the cellular Mg^2+^ content.

The increase in the cellular Mg^2+^ content suppressed the defect not only of 70S ribosome formation, but also the sporulation defect of the Δ*rplA* (L1) mutant (Table 2). Although Spo0A, the master regulator of sporulation, was undetectable in the Δ*rplA* (L1) mutant, Spo0A was clearly produced when the cellular Mg^2+^ content was increased (Fig. 4), and resulted in the restoration of sporulation frequency of the Δ*rplA* (L1) mutant. The phosphorylated form of Spo0A activates expression of sporulation genes as well as its own gene via a positive feedback loop (42). Phosphorylation of Spo0A is achieved by a multicomponent phosphorelay involving at least three kinases called KinA, KinB, and KinC (33, 35). KinA and KinB phosphorylate Spo0F, and phosphorylated Spo0F transfers the phosphoryl group to Spo0B. Finally, Spo0A receives a phosphoryl group from phosphorylated Spo0B (43). In addition, KinC can directly phosphorylate Spo0A without the Spo0F and Spo0B phosphorelay (33). Inversely, phosphorylated Spo0F and Spo0A can be dephosphorylated by phosphatases such as Spo0E (33). It is likely that an inhibition of the multicomponent phosphorelay, and/or the dephosphorylation of Spo0F and Spo0A by the phosphatases was caused by the lack of L1, and it could be suppressed by an increase in the Mg^2+^ content. The improved Spo0A activation in the Δ*rplA* (L1) mutant by Mg^2+^ was probably not due to the complementation of an extraribosomal function of L1, but to restoration of the amount of 70S ribosomes and/or an increase in the cellular translational activity.

In the present study, we demonstrated that the defect in the formation of 70S ribosomes as well as in the sporulation caused by lack of individual ribosomal proteins can be suppressed by increasing the cellular Mg^2+^ content. Mg^2+^ plays a crucial role not only in the ribosome but also in numerous biological processes and cellular functions, such as the activation and catalytic reactions of hundreds of enzymes, utilization of ATP, and maintenance of genomic stability (44, 45). Clarifying the relationship between the ribosome and Mg^2+^, both of which are essential to living cells, is important for understanding cellular function. It has been suggested that the sizes of ribosomal proteins have increased during evolution to complement the function of the rRNA, which originally acted as a ribozyme (46, 47). From another point of view, increasing of the sizes of ribosomal proteins and/or binding of ribosomal proteins to the ribosome during evolution can be considered to complement the Mg^2+^ function in the ribosome, because in the ribosome, the relative abundance of Mg^2+^ is decreased whereas that of ribosomal proteins is increased (47, 48). Further investigation to reveal the mechanism of complementation of the ribosomal-protein function by Mg^2+^ may provide important information about the evolution of the ribosome.

## MATERIALS AND METHODS

### Media and culture conditions

LB medium (49), LB agar, and 2×Schaeffer’s sporulation medium supplemented with 0.1% glucose (2×SG) (50) were used. The culture conditions and media for preparing competent cells have been described previously (51). When required, 5 μg ml^-1^ chloramphenicol, 5 μg ml^-1^ kanamycin and 1 mM isopropyl-β-D-thiogalactopyranoside (IPTG) were added to the media. Growth curves of *B. subtilis* cells were generated by automatically measuring the OD_660_ value of each culture every 5 min using a TVS062CA incubator (ADVANTEC).

### Bacterial strains

All of the *B. subtilis* strains used in this study were isogenic with *B. subtilis* strain 168 *trpC2.* The Δ*rplAv*::*cat*, Δ*rplWv*::*cat*, Δ*rpmJv*::*cat* and Δ*rpsFv*::*cat* strains, which were constructed by replacing the open reading frame of each gene with a promoterless *cat* gene lacking a Rho-independent terminator sequence, were described previously (22). Chromosomal DNA extracted from the Δ*rplAy*::*cat*, Δ*rplWv*::*cat*, Δ*rpmJv*::*cat* and Δ*rpsFv*::*cat* was used to transform the strain harboring Δ*yhdPv*::*erm* and the plasmid pDGmgtE, which carries the *mgtE* gene under the control of an IPTG-inducible Pspac promoter (23), and the transformants were selected on the basis of their chloramphenicol-resistant phenotype.

### Measurement of the cellular Mg^2+^ content

The cellular Mg^2+^ content was measured as described previously (23). Briefly, *B. subtilis* cells were grown in LB medium to exponential phase and harvested. Simultaneously, viable cells were counted by plating the culture on LB agar plates. The cells were resuspended in lysis buffer and disrupted by sonication, and then the pH of the crude extract was adjusted to approximately 3.0 with hydrochloric acid in order to denature the proteins. The amount of Mg^2+^ in the cell lysate was measured with a Metallo Assay Kit for magnesium (Metallogenics). The Mg^2+^ content per cell was calculated by dividing the amount of Mg^2+^ in the crude extract by the number of viable cells. The concentration of Mg^2+^ was calculated by assuming that a *B. subtilis* cell is a cylinder. To measure the cell size (radius and length), microscopic images were analyzed by MicrobeJ, an ImageJ plug-in (52). The mean size of >30 cells was used for the calculation.

### Sucrose density gradient sedimentation analysis

*B. subtilis* cells were grown in LB medium at 37°C with shaking to exponential phase (OD_600_ ~0.4) and harvested. The sucrose density gradient sedimentation analysis was performed as described previously (22). Briefly, the cells were disrupted by passage through a French pressure cell and cell debris was removed by centrifugation. Aliquots of extract were layered onto 10–40% sucrose density gradients, which were subjected to centrifugation at 4°C for 17.5 h at 65,000 ×g (Hitachi P40ST rotor). Samples were collected with a Piston Gradient Fractionator (BioComP), and absorbance profiles were monitored at 254 nm using a Bio-Mini UV Monitor (ATTO, Japan). When normalizing the applied volume by the total absorbance at 260 nm, 10 A_260_ units of crude extract per tube were used.

### Sporulation assay

*B. subtilis* cells were grown in 2× SG medium for 24 h at 37°C with shaking. Heat-resistant spores were counted by heating the cells at 80°C for 10 min, plating them on LB agar plates, and then incubating the plates at 37°C for 24 h.

### Microscopic imaging

*B. subtilis* cells were grown in 2× SG medium at 37°C with shaking. At the indicated times, 500 μl of the culture was removed and subjected to centrifugation at 12,000 × g for 1 min. The cell pellet was resuspended in 40 μl of culture supernatant and then FM4-64 (Invitrogen) and DAPI (Wako Pure Chemical Industries) were added to final concentrations of 10 μg/ml and 5 μg/ml, respectively. The cell suspension was mounted on a microscope slide, coated with poly-L-lysine to fix the cells, and differential interference contrast and fluorescence images were obtained with a LSM800, a confocal fluorescence microscope (Carl Zeiss).

### Western blot analysis

Western blot analysis was performed according to a previously described method (53). Aliquots (15 μg of protein) of crude cell extracts were loaded onto a sodium dodecyl sulfate polyacrylamide gel (12%) and transferred to a PVDF membrane (Millipore Co., Japan). This membrane was then used in the Western blot assay using antisera (1:10,000 dilution) against Spo0A (54).

## ACKNOWLEDGEMENTS

This work was supported in parts by Grants-in-Aid for Scientific Research (C) (26450101 and 15K07013 to G. A. and Y. K.-Y., respectively), Grant-in-Aid for Young Scientists (B) (17K15253 and 23770157 to G. A. and Y. K.-Y., respectively) and Strategic Research Foundation Grant-aided Project for Private Universities (S1201003 to F. K. and Y. K.-Y.) from the Ministry of Education, Culture, Sports, Science, and Technology of Japan.

